# Evidence for a role of synchrony but not common fate in the perception of biological group movements

**DOI:** 10.1101/2023.10.21.563400

**Authors:** Emiel Cracco, Liuba Papeo, Jan R. Wiersema

## Abstract

Extensive research has shown that observers are able to efficiently extract summary information from groups of people. However, little is known about the cues that determine whether multiple people are represented as a social group or as independent individuals. Initial research on this topic has primarily focused on the role of static cues. Here, we instead investigate the role of dynamic cues. In two experiments with male and female human participants, we use EEG frequency tagging to investigate the influence of two fundamental Gestalt principles - synchrony and common fate - on the grouping of biological movements. In Experiment 1, we find that brain responses coupled to four point-light figures walking together are enhanced when they move in sync vs. out of sync, but only when they are presented upright. In contrast, we found no effect of movement direction (i.e., common fate). In Experiment 2, we rule out that synchrony takes precedence over common fate by replicating the null effect of movement direction while keeping synchrony. These results put forward synchrony as an important driver of social grouping, consistent with the fact that it is an important feature of social interaction and an indicator of social cohesion. In contrast, the influence of common fate on social grouping is less clear and will require further research.

## Introduction

Social interaction is at the heart of human life: much of what we do, we do with others. Understanding social interactions requires processing not only individual actions, but also how they relate to one another. Although the processing of biological movements (e.g., Blake & Shiffrar, 2007; Caspers et al., 2010; Pitcher & Ungerleider, 2021) and how this scales up from one to multiple agents (e.g., Cracco et al., 2015, 2016, 2019, 2022; Cracco & Brass, 2018) is well understood, relatively little is known about how we organize agents into groups.

What is known, is that observers can efficiently extract summary information from groups of people (e.g., Nguyen et al., 2021; Sweeny et al., 2013; Sweeny & Whitney, 2014; Whitney & Leib, 2018). However, an important open question is what determines whether multiple agents are seen as a group, or as rather as multiple individuals (Hafri & Firestone, 2021; Kaiser et al., 2019; Papeo, 2020). Initial research into this question has focused on static cues and found that individuals who appear to interact by virtue of their relative spatial positioning (e.g., standing face-to-face) are grouped in working memory (e.g., Ding et al., 2017; Paparella & Papeo, 2022; Vestner et al., 2019) and visual perception (e.g., Adibpour et al., 2021; Bellot et al., 2021; Ji et al., 2020; Papeo, 2020; Papeo et al., 2017). Here, we instead investigate two movement cues: synchrony (i.e., temporal alignment) and common fate (i.e., spatial alignment). Both synchrony and common fate are important Gestalt principles that play a fundamental role in structuring basic motion input into coherent percepts (for reviews, see Wagemans, Elder, et al., 2012; Wagemans, Feldman, et al., 2012). Yet, whether they also contribute to organizing more complex scenes with multiple social agents remains to be explored.

Recent research has provided initial evidence that synchrony can indeed shape the processing of other people’s actions (Alp et al., 2017; Cheng et al., 2022; Cracco, Lee, et al., 2022; Tsantani et al., 2022) or emotional expressions (Elias et al., 2017). However, in addition to being restricted to synchrony, this research has not yet established whether these effects are specific to biological agents or are more general in nature. Here, we address this question by simultaneously investigating how synchrony and common fate influence the perception of biological group movements. To do so, we adapted a recently developed frequency tagging task that measures the holistic processing of biological movement (Cracco, Oomen, et al., 2022). Specifically, we extracted brain responses coupled to the walking pace of 4 point-light figures moving at a fixed pace either in or out of synchrony (synchrony), in the same or different directions (common fate), upright or inverted (inversion). Experiment 1 manipulated both synchrony and common fate. Experiment 2 manipulated common fate alone.

If synchrony and common fate contribute to social grouping, brain responses at the figures’ walking pace should be modulated by these principles. Previous research has shown that neural activity in visual brain areas is increased when objects (Kim & Biederman, 2011; Roberts & Humphreys, 2010) or bodies (Abassi & Papeo, 2020, 2022) are organized in interactive vs. non-interactive constellations (e.g., facing), and these effects were found to correlate with behavioral indices of perceptual grouping (Abassi & Papeo, 2022). As a result, processing multiple walkers as a single group should result in stronger brain responses coupled to those movements. If both cues influence perception through the same mechanism, we might furthermore expect them to interact (e.g., Han, 2004; Lee & Blake, 2001; Peterson & Kimchi, 2013).

In addition, if synchrony and common fate influence biological motion perception, rather than motion perception more generally, we should see that their effects are sensitive to inversion. Indeed, inversion is well-known to disrupt the configural processing of biological motion (Cracco, Oomen, et al., 2022; Giese & Poggio, 2003) in that it triggers a local processing style where point-light figures are processed as local dots instead of global agents (Bertenthal & Pinto, 1994; Pavlova & Sokolov, 2000). Therefore, if synchrony and common fate influence social grouping, as opposed to a more unspecific grouping of objects moving in synchrony or in the same direction, their effects should be abolished when the walkers are inverted.

## Materials and Methods

### Open Science Statement

Both experiments were preregistered (https://aspredicted.org/NRY_639 and https://aspredicted.org/SQM_V4G). The stimuli, data files, analysis script, and experimental program are available on the OSF together with example videos (https://osf.io/67sn8/).

### Experiment 1

#### Participants

In line with our preregistration, we collected a sample of 33 volunteers who participated in the experiment in return for €25. Individuals could only participate if they were between 18 and 35 years old, had normal or corrected-to-normal vision, were right-handed, were fluent in Dutch, and did not have a history of neurological or psychiatric disorder. Age restrictions were imposed to ensure a homogenous sample and limit age-related effects. The sample size was based on a previous study in which we found a medium-sized effect (*d*_z_ = 0.56) of inversion (Cracco, Oomen, et al., 2022). Specifically, we conservatively rounded down this effect to *d*_z_ = 0.50 and calculated the sample size needed to obtain 80% power to detect such an effect at α = 0.05. This revealed a required sample size of N = 33. Following our preregistered approach, two participants were excluded due to bad data quality (> 10% of electrodes requiring interpolation^1^), resulting in a final sample of 31 participants (24 female, 7 male, *M*_age_ = 22.90, range_age_ = 18-32). The study was approved by the local ethics board of the Faculty of Psychological and Educational Sciences at Ghent University (2021/129).

#### Task, Stimuli, and Procedure

Participants first signed the informed consent form. Next, they filled out the Dutch version of the Autism Quotient questionnaire (AQ; Hoekstra et al., 2008) while the experimenter prepared the EEG cap. The AQ was included for exploratory purposes, based on evidence that autism is associated with a decreased ability or propensity to extract socially relevant information from biological motion (Federici et al., 2020; Todorova et al., 2019). Internal consistency of the AQ in the current study was good (α = 0.78).

After preparing the EEG cap, the experiment was started. The experiment was programmed in PsychoPy (Peirce et al., 2019) and rendered on a 24-inch computer monitor with a 60 Hz refresh rate, positioned approximately 80–100 cm from the participant. During the experiment, participants watched videos of 4 point-light walkers, walking in place at a fixed pace of 2.4 Hz (1 step every ∼417 ms). Each walker extended an area of ∼2.33° x 5.24°, with a horizontal distance of 4.08° between the mid-point of two walkers and a vertical distance of 6.69°. In a previous study, we found that a walking pace of 2.4 Hz elicits a brain response at the frequency of walking and that this response disentangles biological motion perception from the perception of local motion or other lower-level features of the stimuli (Cracco, Oomen, et al., 2022). The walkers (2 male, 2 female) were created with the online <u>BMLStimuli</u> tool (Troje, 2002, 2008) and were organized symmetrically around a salmon-colored fixation cross (see Figure 1). The locations of the four walkers in the display were counterbalanced across participants, with the restriction that neighboring figures on both sides were never of the same sex to avoid horizontal or vertical grouping based on similarity.

**Figure 1.**
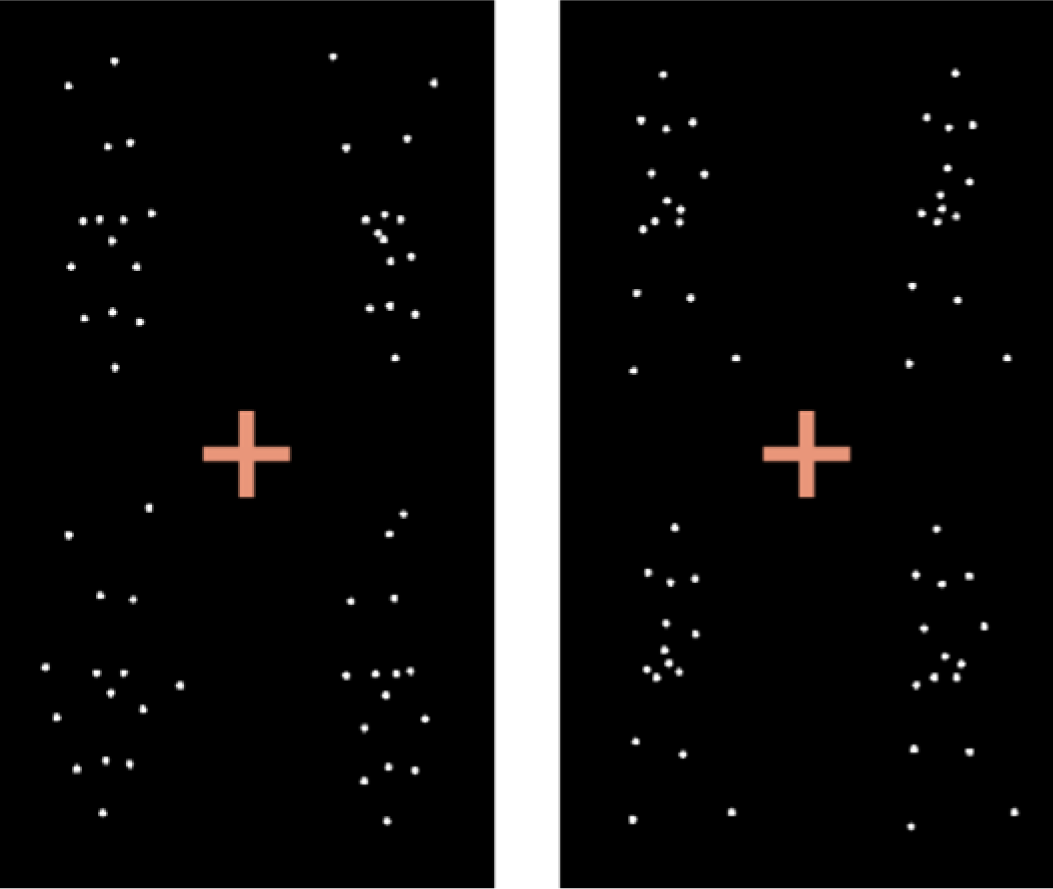
Still frames from two example stimuli used in Experiment 1. The left example shows four inverted walkers. The right example shows four upright walkers. Example videos of each condition can be found on the Open Science Framework (OSF; https://osf.io/67sn8/).

Each participant watched 32 videos in total. Each video lasted exactly 124 walker steps (∼52 s), including a 4-step fade-in and a 4-step fade-out period (∼2 s each). Relatively long video durations were used to obtain a high spectral resolution (= 1/video duration). The 32 videos were randomly assigned to one condition of the movement direction (same vs. different) x synchrony (synchronous vs. asynchronous) x orientation (upright vs. inverted) design. Each of these 8 conditions was repeated 4 times. To minimize habituation, the walking direction of the 4 walkers differed between the different repetitions of the same condition.

In the *same direction* conditions, the azimuth of the 4 walkers (i.e., the rotation of the walker around the vertical axis) was either -135°, -45°, 45°, or 135°. In the *different direction* conditions, each walker was assigned one of those azimuths, with the restriction that walkers were never oriented toward each other and were never oriented to create an X-like pattern where, expressed in 2D terms, the top-left agent walked to the top-left corner, the top right agent to the top right corner, etc. This was done to prevent that a clear structure emerged from the way in which the walkers moved. Fixed azimuths were used to prevent accidental facingness or other social constellations (e.g., walking toward another walker) in the *different direction* condition. Importantly, to ensure that the azimuths were associated with unambiguous percepts, point-light stimuli were rendered using perspective projection (Vanrie et al., 2004).

In the *synchrony* conditions, all walkers started from the same position in the walking cycle. In the *asynchrony* conditions, all walkers started from different (random) positions in the walking cycle, which were at least 10 frames apart from each other. The start positions stayed the same across the experiment but differed across participants. All above stimuli were presented both upright and inverted. To maintain attention and control eye gaze, participants were instructed to look at the fixation cross around which the walkers were organized and to press the space bar every time it briefly (400 ms) turned red. This happened 2 to 4 times per video, at random time points. Accuracy of participants on this task was high and stable across the eight conditions (93–95%).

#### Rating Task

At the end of the experiment, participants performed a rating task in which they saw shortened videos (10s) of the same stimuli and had to answer the following questions in random order after each video: (i) Did the figures move in the same direction? (ii) How synchronous did the figures move?, (iii) How pleasant did you find the video to look at?, (iv) How complex did you find the video? This rating task had two purposes. First, it was included to test if participants could detect whether the walkers moved in or out of synchrony and whether they moved in the same or in different directions. Second, it was included as a replication of Cracco, Lee, et al. (2022), who found that synchronous videos were perceived as less complex and more aesthetically pleasing than non-synchronous videos. In contrast to the main task, each condition of the direction x synchrony x orientation design was shown only once in the rating task. The direction question was answered by clicking a ‘yes’ or ‘no’ button. The other three questions were answered by moving a cursor along a scale from 0 to 100.

#### EEG Preprocessing

EEG was recorded using an ActiCHamp amplifier and BrainVisionRecorder software (version 1.21.0402, Brain Products, Gilching, Germany) at a sampling rate of 1000 Hz with 64 active Ag/AgCI electrodes. Electrode positions were based on the 10%-system, except that TP9 and TP10 were replaced with OI1h and OI2h according to the 5%-system to better cover posterior scalp sites. Fpz was used as the ground electrode and Fz as the online reference. FT9 and FT10 were used to record horizontal eye movements and two Ag/AgCI sintered ring electrodes placed above and below the left eye to record vertical eye movements.

The EEG signal was processed offline in Letswave 6 by band-pass filtering the raw data between 0.1 Hz and 100 Hz, segmenting the filtered data into 8 experimental conditions, and applying ICA (RUNICA algorithm, square mixing matrix) to the merged segmented data. The first 10 components were inspected, and those capturing eye blinks or horizontal eye movements were removed. Faulty or excessively noisy electrodes were interpolated from the 3 closest neighbors (1% on average, never more than 10%). Fz was then reinserted, and the data were re-referenced to the average signal across all electrodes. To ensure that the epoch length was a multiple of the presentation rate, epochs were cropped from the end of the fade-in to the start of the fade-out period. Finally, conditions were averaged, and the discrete Fourier transform of the signal was computed using a fast Fourier transform algorithm, resulting in normalized amplitudes (μV) in the frequency domain with a bin size of ∼0.02 Hz (∼1/48.33).

#### Data Analysis

As frequency tagging elicits responses not only at the tagged frequency (2.4 Hz) but also at harmonics of that frequency (4.8 Hz, 7.2 Hz, …), we quantified the brain response as the sum of the baseline-subtracted amplitudes across all relevant harmonics (for further justification, see Retter et al., 2021; Retter & Rossion, 2016). As preregistered, and following Cracco, Oomen, et al. (2022), we identified the relevant harmonics by (i) calculating the grand-averaged amplitudes across participants, conditions, and electrodes, (ii) transforming the amplitudes at each frequency bin into z-scores, using as baseline the 10 surrounding bins on each side, excluding directly adjacent bins, and (iii) identifying the harmonics with a z-score > 2.32 (i.e., p < .01). This revealed 2 relevant harmonics, at 2.4 Hz and 4.8 Hz. For these two harmonics, we then calculated baseline-subtracted amplitudes using the same baseline as before and summed them to quantify the brain response (Figure 2).

**Figure 2.**
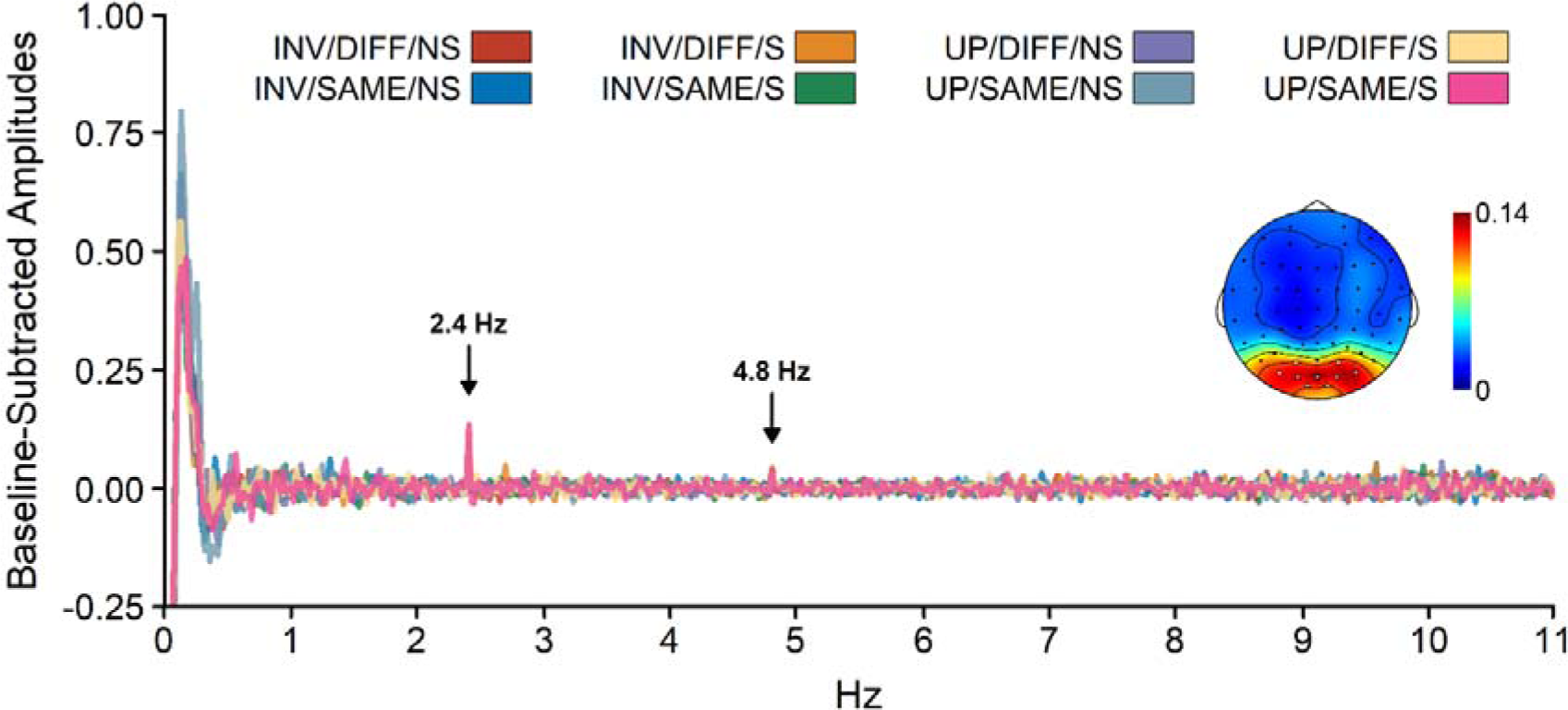
Collapsed topography and spectrum plot of Experiment 1. The collapsed topography shows the sum of the baseline-subtracted amplitudes at 2.4 and 4.8 Hz, averaged across participants and conditions. Electrodes included in the analysis are marked in white. The topography is scaled from 0 to the maximum across electrodes (0.14 μV). The spectrum plot shows the baseline-subtracted amplitudes in the 8 conditions, averaged across participants and the electrodes included in the statistical analysis. INV: inverted, UP: upright, DIFF: different directions, SAME: same direction, NS: Non-synchronous, S: Synchronous.

In addition to a response at 2.4 Hz, we also preregistered to explore the presence of a response at 1.2 Hz, which is the frequency at which each individual dot repeats its trajectory. In a previous study, we had found responses at this frequency, but only for scrambled walkers (Cracco, Oomen, et al., 2022). In line with this finding, amplitudes at 1.2 Hz (and harmonics) did not exceed the predefined z-score threshold of 2.32. As a result, these frequencies were not considered further.

To select the electrodes to include into the analysis, we used a collapsed localizer approach (Luck & Gaspelin, 2017). More precisely, we averaged the summed baseline-subtracted amplitudes across participants and conditions and chose the electrodes to include from the collapsed topography. This topography revealed activity over occipital and parieto-occipital electrodes. Based on this, we decided to include three electrode clusters in the analysis: left posterior (PO7, PO3, O1), middle posterior (OI1h, Oz, OI2h), and right posterior (O2, PO4, PO8). More specifically, we analyzed the brain response with a cluster (left vs. middle vs. right) x movement direction (same vs. different) x synchrony (synchronous vs. asynchronous) x orientation (upright vs. inverted) repeated measures ANOVA. In addition, as a preregistered exploratory analysis, we also correlated the brain response with the collected AQ scores. The rating data were analyzed using separate movement direction (same vs. different) x synchrony (synchronous vs. asynchronous) x orientation (upright vs. inverted) repeated measures ANOVAs for the four ratings (synchrony, direction, complexity, and liking).

### Experiment 2

#### Participants

Following our preregistration, we collected a new sample of 33 volunteers using the same procedure as in Experiment 1. However, one participant had to be excluded due to bad data quality (> 10% of electrodes requiring interpolation), resulting in a final sample of 32 participants (27 female, 5 male, *M*_age_ = 22.16, *range*_age_ = 17-32).

#### Task, Stimuli, and Procedure

All procedures were identical to Experiment 1, except that Experiment 2 only included videos where the walkers moved synchronously and no longer included the rating task at the end. The internal consistency of the AQ was good (α = 0.92). Accuracy of participants on the attention check was 97–98% in all conditions.

#### EEG Preprocessing and Data Analysis

The preprocessing and analysis pipelines were identical to Experiment 1. In line with Experiment 1, only the first two harmonics (2.4 and 4.8 Hz) reached our predefined z-score cut-off (Figure 3) and the baseline-subtracted amplitudes at these two frequencies were summed to quantify the brain response. The collapsed localizer of Experiment 2 revealed only a right posterior cluster. Though inconsistent with the collapsed topography of Experiment 1, such a right-lateralized scalp pattern matches the topography of the synchrony conditions in Experiment 1. For consistency, we therefore included not just the right posterior cluster (O2, PO4, PO8), but also the left (PO7, PO3, O1) and middle posterior (OI1h, Oz, OI2h) clusters in the analysis of Experiment 2. The resulting brain responses were analyzed with a cluster (left vs. middle vs. right) x movement direction (same vs. different) x orientation (upright vs. inverted) repeated measures ANOVA.

**Figure 3.**
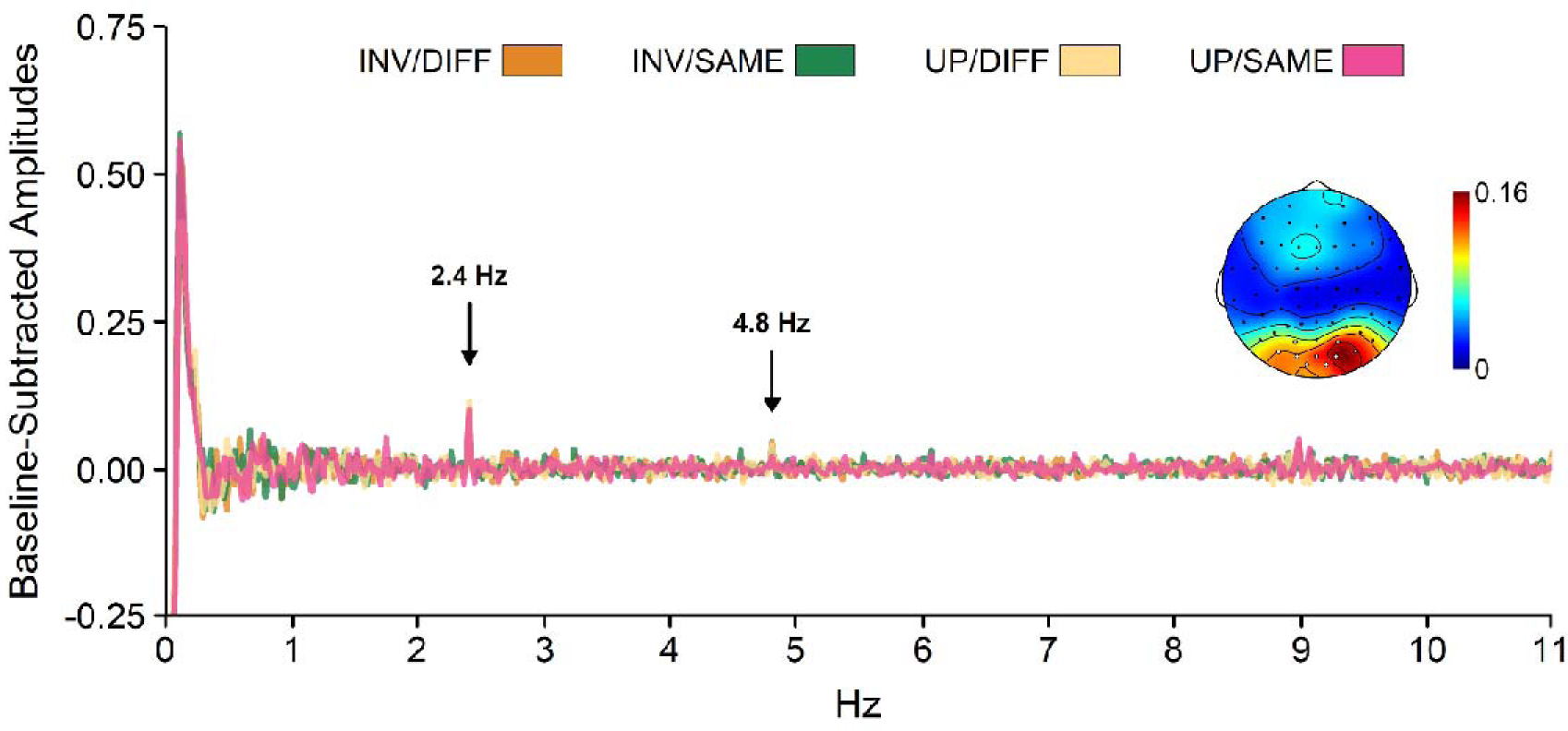
Collapsed topography and spectrum plot of Experiment 2. The collapsed topography shows the sum of the baseline-subtracted amplitudes at 2.4 and 4.8 Hz, averaged across participants and conditions. Electrodes included in the analysis are marked in white. The topography is scaled from 0 to the maximum across electrodes (0.16 μV). The spectrum plot shows the baseline-subtracted amplitudes in the 4 conditions, across participants and the included electrodes. INV: inverted, UP: upright, DIFF: different directions, SAME: same direction.

## Results

### Rating Task

Due to an oversight, the rating task was not administered to one participant. Hence, analyses were conducted on 30 out of 31 participants in Experiment 1. The rating task had two aims. First, we sought to confirm that our participants could differentiate between walkers moving in or out of synchrony and between walkers moving in the same or in different directions. Second, we sought to replicate a previous finding that groups moving in synchrony are perceived as less complex and as more aesthetically pleasing (Cracco, Lee, et al., 2022). As such, we will focus on the contrasts answering those questions. However, a full overview of all rating results can be found on the OSF (https://osf.io/67sn8/).

With respect to our first aim, we found significant effects of synchrony on the synchrony ratings, *F*(1, 29) = 41.91, *p* < .001, η^2^_p_ = 0.59, and of movement direction on the movement direction ratings, *F*(1, 29) = 333.39, *p* < .001, η^2^_p_ = 0.92. The synchrony effect indicated that participants rated the walkers as more synchronous when they moved in synchrony (*M* = 75, *SD* = 20) than out of synchrony (*M* = 49, *SD* = 18). The movement direction effect indicated that participants were more likely to rate the walkers as going in the same direction when they indeed did so (*M* = 91%, *SD* = 15%), as compared to when they did not (*M* = 17%, *SD* = 19%).

With respect to our second aim, we found that both complexity ratings, *F*(1, 29) = 16.74, *p* < .001, η^2^_p_ = 0.37, and liking ratings, *F*(1, 29) = 34.58, *p* < .001, η^2^_p_ = 0.54, were influenced by synchrony. Whereas complexity was rated lower when the walkers moved in sync (*M* = 37, *SD* = 14) then when they moved out of sync (*M* = 49, *SD* = 16), liking was rated higher for synchronous (*M* = 63, *SD* = 16) than for asynchronous walkers (M = 46, SD = 15).

### Effects of Synchrony and Common Fate on Movement Processing

To investigate the effects of synchrony and common fate on the processing of biological group movements, Experiment 1 investigated the influence of both variables on the EEG response coupled to the walking frequency of the four point-light agents. As shown in Figure 4, this revealed a main effect of synchrony, *F*(1, 30) = 8.86, *p* = .006, η^2^_p_ = 0.23, with larger amplitudes for synchronous than for asynchronous walkers, but no effect of movement direction, *F*(1, 30) = 3.07, *p* = .090, η^2^_p_ = 0.09. Though not statistically significant, the effect of movement direction was opposite to what we predicted, with slightly larger amplitudes when the walkers moved in different directions than when they moved in the same direction. In addition to the main effect of synchrony, we also found a cluster x synchrony interaction, *F*(2, 29) = 7.41, *p* = .003, η^2^_p_ = 0.34, and a synchrony x orientation interaction, *F*(1, 30) = 9.88, *p* = .004, η^2^_p_ = 0.25. The cluster x synchrony interaction indicated that there was a synchrony effect in the middle, *t*(30) = 2.98, *p* = .006, *d*_z_ = 0.54, and right clusters, t(30) = 3.79, p < .001, dz = 0.68, but not in the left cluster, *t*(30) = 1.08, *p* = .291, *d*_z_ = 0.19. The synchrony x orientation interaction indicated that there was a synchrony effect for upright, *t*(30) = 4.57, *p* < .001, *d*_z_ = 0.82, but not for inverted walkers, *t*(30) = 0.48, *p* = .637, *d*_z_ = 0.09. None of the other effects reached significance, all *ps* ≥ .071. Finally, exploratory correlations with the AQ total scores revealed that neither the main effect of synchrony, *r* = .12, *p* = .507, nor the synchrony x orientation interaction, *r* = .25, *p* = .170, correlated significantly with autistic traits.

**Figure 4.**
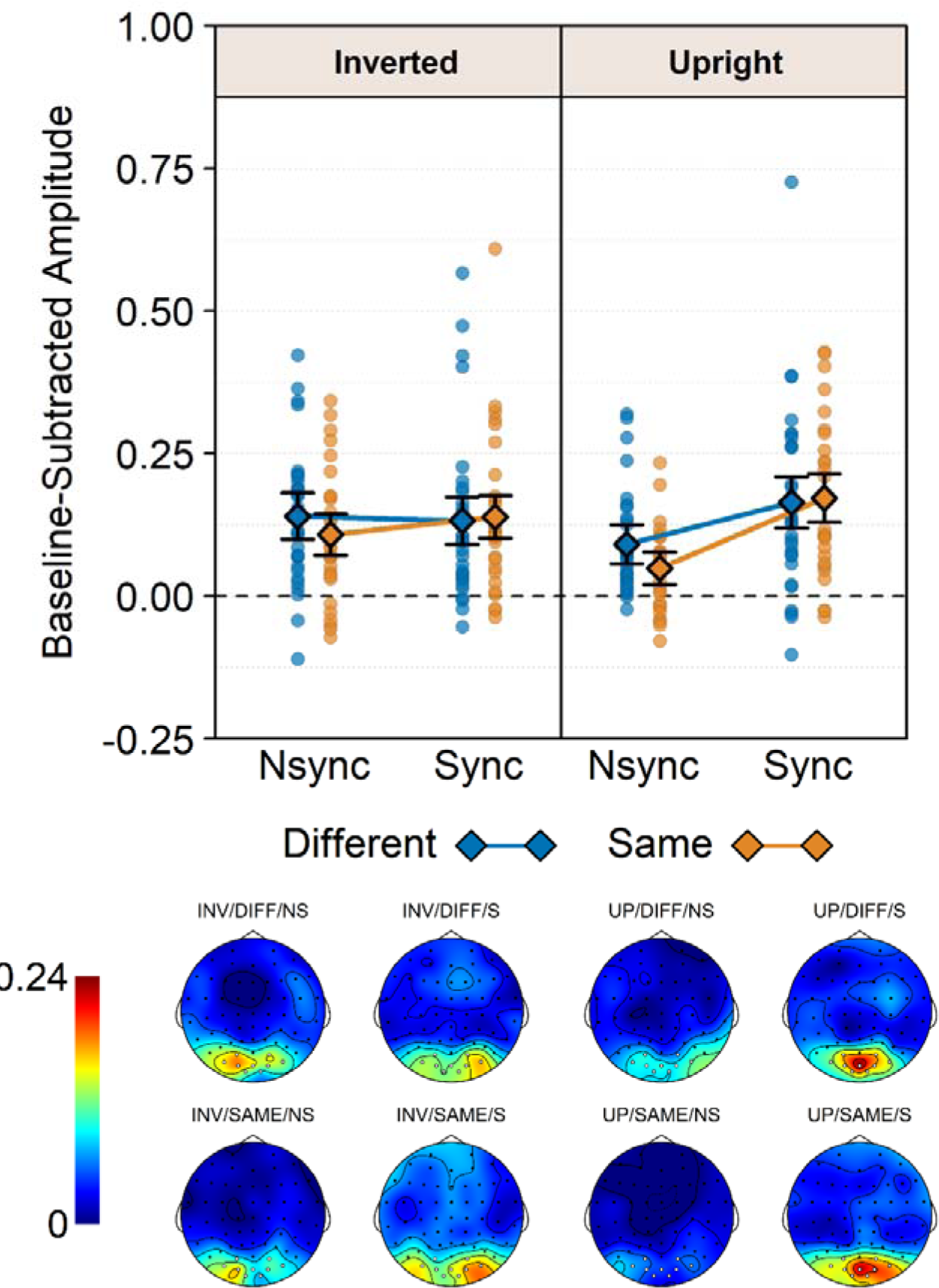
Baseline-subtracted amplitudes at the walking frequency (2.4 Hz) and its harmonics in Experiment 1. Note that 0 is the baseline and hence that values below 0 reflect noise. The dots show the amplitude of the brain signal per condition and per participant. The diamonds show the amplitudes per condition across participants. Error bars are 95% confidence intervals corrected for within-subject designs (Morey, 2008). Topographies are scaled from 0 to the maximum across the scalp and conditions (0.24 μV). INV: inverted, UP: upright, DIFF: different directions, SAME: same direction, NS: Non-synchronous, S: Synchronous.

These results show that synchrony influenced the neural processing of biological group movements, while common fate did not. Importantly, this is true even though the results of the rating task showed that participants could reliably detect whether the walkers moved in the same or different directions. Our results therefore suggest that the movement synchrony but not common movement direction is used to organize individuals in groups. However, given that it is still unclear whether and how the brain integrates Gestalt cues (Peterson & Kimchi, 2013), an alternative explanation could be that differences in synchrony take precedence over differences in common fate when organizing stimuli into groups. In support of this idea, we have previously found that image sequences eliciting fluent (i.e., spatiotemporally consistent) movement yield stronger brain responses than those eliciting non-fluent movement when there is only one agent (Cracco, Lee, et al., 2022) or four agents who always move in synchrony (Cracco et al., 2023), but not when movement fluency is manipulated together with synchrony (Cracco, Lee, et al., 2022). It is also consistent with evidence that individuals moving in the same direction are rated high on social cohesion, but not when they move asynchronously (Wilson & Gos, 2019). In Experiment 2, we therefore investigated whether an effect of common fate would emerge when synchrony was kept constant.

### Effect of Common Fate on Movement Processing When Synchrony is Constant

Experiment 2 tested whether common fate would influence the processing of biological group movements when the walkers always moved in synchrony. As shown in Figure 5, this was not the case: movement direction did not influence the EEG response coupled to the walking pace of the four point-light agents, *F*(1, 31) = 2.99, *p* = .094, η^2^_p_ = 0.09. If anything, the effect again went in the opposite direction relative to what we predicted, with a stronger response to agents moving in different directions *vs.* in the same direction. The effect of cluster was non-significant, *F*(2, 30) = 2.63, *p* = .089, η^2^_p_ = 0.15, but showed a trend for a right lateralized effect with larger response amplitudes in the right cluster than in the middle cluster and larger amplitudes in the middle cluster than in the left cluster. None of the remaining main effects or interactions were significant, all *ps* ≥ .180. Exploratory correlations with the AQ total scores revealed that neither the main effect of movement direction, *r* = .02, *p* = .933, nor the movement direction x orientation interaction, *r* = .07, *p* = .710, correlated with autistic traits. These results eliminate the possibility that the absence of a common fate effect in Experiment 1 was due to differences in synchrony taking precedence over differences in common fate. Instead, they suggest that only synchrony contributes to social grouping.

**Figure 5.**
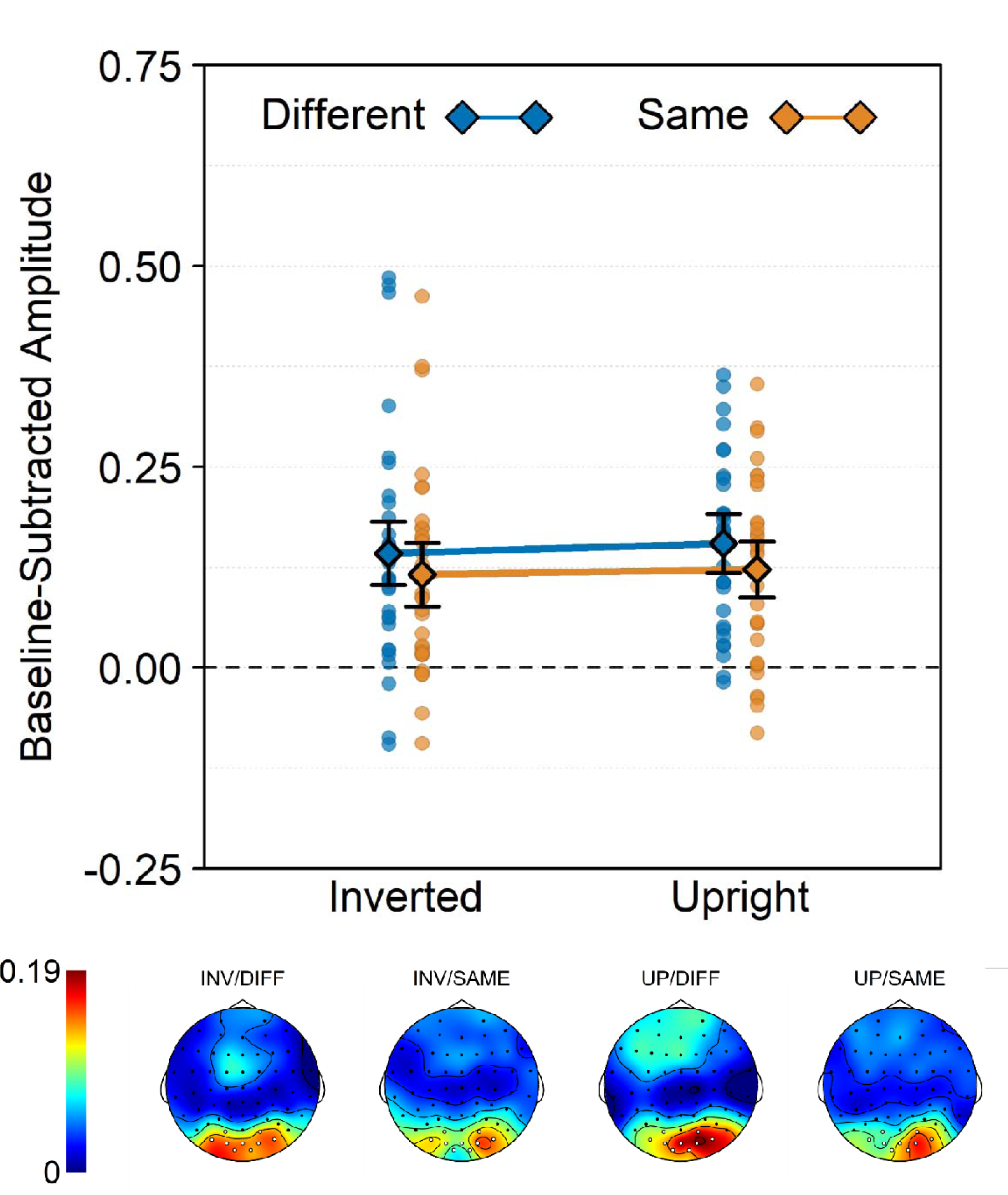
Baseline-subtracted amplitudes at the walking frequency (2.4 Hz) and its harmonics in Experiment 2. As 0 is the baseline, values below 0 reflect noise. The dots show amplitudes per condition and per participant. The diamonds show amplitudes per condition across participants. Error bars are 95% confidence intervals corrected for within-subject designs (Morey, 2008). Topographies are scaled from 0 to the maximum across the scalp and conditions (0.19 μV). INV: inverted, UP: upright, DIFF: different directions, SAME: same direction.

## Discussion

Observers are able to efficiently extract summary information from groups of people (e.g., Nguyen et al., 2021; Sweeny et al., 2013; Sweeny & Whitney, 2014; Whitney & Leib, 2018). However, not much is known about the cues that determine whether multiple individuals are represented as independent agents or as a group. In particular, even though motion is essential to social perception (Pitcher & Ungerleider, 2021), most research so far has studied static cues contributing to social grouping (e.g., facing each other; Papeo, 2020). Here, we studied the role of dynamic cues instead. Inspired by Gestalt perception, we investigated in two experiments whether two well-known Gestalt principles – synchrony and common fate – influence the processing of four point-light figures walking in proximity to each other. As a marker of social grouping, we analyzed the extent to which a frequency-tagged EEG response, previously associated with the holistic perception of biological motion (Cracco, Oomen, et al., 2022), was facilitated when the walkers’ movements could be organized into a fluent, unified percept according to synchrony and/or common fate. The results revealed that brain responses coupled to the walkers’ movements were facilitated when the walkers moved in synchrony, but only when they were shown upright (Experiment 1). In contrast, common fate did not reliably alter the brain response, neither when it was manipulated together with synchrony (Experiment 1), nor when it was manipulated alone (Experiment 2).

### The Role of Synchrony in Biological Motion Perception

Synchrony between the movements of upright walkers resulted in stronger brain responses. The finding that brain activity was enhanced is consistent with evidence that brain activity in visual brain areas is increased when objects (Kim & Biederman, 2011; Roberts & Humphreys, 2010) or bodies (Abassi & Papeo, 2020, 2022) can be represented as a single unit. The finding that this effect was restricted to upright walkers further indicates that it was driven by global, not local synchrony. Indeed, when using the same task with a single agent, we found that inversion specifically disrupted global processing of the walker’s movements, while leaving processing of the local dot trajectories untouched (Cracco, Oomen, et al., 2022). Together, our results thus indicate that synchrony influenced grouping at the level of the social agents.

These results add onto a burgeoning literature investigating synchrony in social perception (Alp et al., 2017; Cheng et al., 2022; Cracco, Lee, et al., 2022; Tsantani et al., 2022). In a functional MRI study, Tsantani et al. (2022) showed that synchrony between the head movements of two agents could be decoded in brain areas of the social perception network, such as the posterior superior temporal cortex and the extrastriate body area.

However, this study did not include a basic motion control, leaving open the question of whether synchrony influences social perception or motion perception more generally. Using EEG frequency-tagging, Alp et al. (2017) and Cracco et al. (2022) did include such a control in the form of inverted stimuli, a condition known to disrupt body representation (Reed et al., 2003). However, in contrast to the current study, where synchrony only affected perception of upright stimuli, those previous studies found independent effects of synchrony and inversion: stronger brain responses for upright vs. inverted stimuli and for synchronous vs. asynchronous stimuli.

The differences between previous work and the current study may be related to the measured brain response. Like the present work, both Alp et al. (2017) and Cracco et al. (2022) used frequency tagging to measure the response to biological motion perception.

However, Alp et al. (2017) modulated the contrast of four point-light agents dancing in or out of synchrony and measured the brain responses coupled to the frequency of contrast modulation. As this frequency was independent of the movements performed by the agents, this means that they did not tag the movements themselves, but rather the visual appearance of the agents performing those movements. Cracco et al. (2022) did tag the movements themselves, but measured the top-down reconstruction of those movements from static input, rather than their bottom-up processing from motion kinematics. To be more precise, Cracco et al. (2022) did not show dynamic stimuli but instead showed sequences of static images.

Within a range of inter-stimulus-intervals, such sequences can elicit an apparent movement percept in the absence of retinal motion (e.g., Grosjean et al., 2009; Orgs et al., 2011; Shiffrar & Freyd, 1990). However, this type of motion perception is thought to rely on different neural processes (Giese & Poggio, 2003) and recruits a different brain network than the network involved in perceiving biological motion from kinematics (Orgs et al., 2016; Stevens et al., 2006).

In line with the fact that synchrony describes the dynamic relationship between two or more movement paths, a comparison of the current work with earlier research thus indicates that synchrony may specifically interact with the processing of biological movements from motion kinematics, rather than the processing of the agent performing those movements (Alp et al., 2017) or the top-down reconstruction of movements from static input (Cracco, Lee, et al., 2022).

### The Role of Common Fate in Social Grouping

In contrast to synchrony, we found no evidence that common fate, another well-known Gestalt principle, influenced biological motion perception. There are several possible explanations for this finding. First, it could be that participants had difficulties to distinguish between walkers moving in the same vs. different directions. However, this seems unlikely based on the rating task results: participants could clearly differentiate between walkers based on whether they moved in one or in multiple directions. Importantly, while error rates were relatively high when the walkers moved in multiple directions (17%), a closer look at the data split up for the different conditions revealed that this was entirely driven by errors in the conditions where the walkers were inverted (34%), whereas not a single error was made when they were shown upright (https://osf.io/67sn8/). This is consistent with previous work showing that observers are highly accurate at differentiating between the walking directions of point-light figures, at least when perspective projection is used, like in the current study (Vanrie et al., 2004).

Another possible reason for why synchrony but not common fate influenced the brain response is that they operated at different levels. Indeed, albeit non-significant, both experiments found slightly *stronger* responses when the walkers moved in different directions than when they moved in the same direction. Though speculative, one interpretation for this pattern could be that common fate influences the processing of the individual dot movements (i.e. local processing of biological motion) and therefore that seeing multiple walkers move in the same direction increases focus on the individual dots while decreasing focus on the point-light agents (i.e., global processing of biological motion). This is consistent with evidence that motion coherency in random dot motion paradigms influences brain activity in brain areas typically associated with local processing of biological motion (e.g., V5/MT; Braddick et al., 2001; Rees et al., 2000; Rina et al., 2022), whereas the brain response measured here has been shown to specifically capture global processing of biological motion (Cracco, Oomen, et al., 2022).

The absence of a strong common effect could also be related to the method. More specifically, it is possible that EEG frequency tagging is inherently more sensitive to effects of synchrony. Indeed, whereas the degree of synchrony between brain waves directly influences the amplitude of the combined brain response, there is no such direct connection for common fate. As such, it is possible that the imbalance between synchrony and common fate here was caused by the method and that other methods may find an influence of common fate on social grouping. An interesting technique in this respect would be functional MRI, which has been successfully used to demonstrate effects of common fate on the processing of more basic, non-social stimuli (e.g., Braddick et al., 2001; Rees et al., 2000; Rina et al., 2022).

A fourth possible reason for the absence of a common fate effect is that common fate may be less relevant for social perception and for that reason did not influence the brain response. While the social relevance of synchrony has long been acknowledged, based on the fact that it is a common factor to many social interactions (Thurman & Lu, 2014) and a consistent indicator of social cohesion (Lakens, 2010; Lakens & Stel, 2011; Wilson & Gos, 2019), the social function of common fate is less clear. On the one hand, one could argue that a common movement direction is indicative of a common goal and therefore of social cohesion. On the other hand, there is currently little evidence that common fate in and of itself influences social perception. To our knowledge, there is only one study so far that has investigated whether observers ascribe different social dynamics to groups moving in the same vs. different directions (Wilson & Mansour, 2020). However, ‘collective movement’ in that study was operationalized as moving synchronously in the same direction, in close proximity of each other. In other words, collective movement in Wilson & Manour (2020) combined common fate with synchrony and proximity. Moreover, when participants were presented with two dots that always moved in the same direction, but did so asynchronously and without maintaining close distance, social cohesion was rated only 2.36 out of 7 (Wilson & Mansour, 2020). While more research is needed, this suggests a potential imbalance between synchrony and common fate, with the former but not the latter having an important social function.

A directional cue that is potentially more relevant for social grouping than common fate is whether individuals move towards or away from each other. In line with this idea, recent work has shown that people facing towards each other are detected more easily (Papeo et al., 2017, 2019; Vestner et al., 2019) and remembered better (Paparella & Papeo, 2022; Vestner et al., 2019) than people facing away from each other. In line with the synchrony effects found here, these effects were sensitive to inversion (Papeo et al., 2017, 2019; Vestner et al., 2019) and were visible in the brain as increased integrative response (Adibpour et al., 2021) and enhanced activity in body-(Abassi & Papeo, 2020) and movement-related brain areas (Bellot et al., 2021). Although there is an ongoing discussion as to whether these effects are social in nature (Papeo, 2020) or reflect a more general attentional mechanism (Vestner et al., 2020, 2021), they suggest that organizing people into groups may depend not on whether they move in the same or in different directions, but on whether they move towards or away from each other.

## Conclusion

The current study shows that social grouping is driven not only by static (e.g., facing each other; Papeo, 2020), but also by dynamic cues. Inspired by Gestalt perception (Wagemans, Elder, et al., 2012; Wagemans, Feldman, et al., 2012), we investigated the role of synchrony and common fate. The results revealed that synchrony influenced the perception of biological group movements, whereas common fate did not. An important question for future research will be to investigate whether such effects can be observed with other methods and/or stimuli, or that there is a more fundamental disparity between synchrony and common fate, perhaps related to differences in the extent to which they signal social cohesion or social interaction.

## Ethics statement

The study was approved by the local ethics board of the Faculty of Psychological and Educational Sciences at Ghent University (2021/129).

## Data availability statement

The stimuli, data files, analysis script, and experimental program are available on the OSF (https://osf.io/67sn8/).

## Acknowledgements

This work was funded by a senior postdoctoral fellowship awarded to EC by the Research Foundation Flanders (12U0322N). LP was supported by a European Research Council Grant (Project THEMPO-758473).

## Footnotes

1 Note that this was an arbitrary threshold. However, it was preregistered and in line our previous work (Cracco et al., 2023; Cracco, Oomen, et al., 2022).

